# *In vitro* effects of S-Licarbazepine as a potential precision therapy on *SCN8A* variants causing neuropsychiatric disorders

**DOI:** 10.1101/2021.04.24.441205

**Authors:** Erva Bayraktar, Yuanyuan Liu, Ulrike B.S. Hedrich, Yildirim Sara, Holger Lerche, Thomas V Wuttke, Stephan Lauxmann

## Abstract

**Background and Purpose:** Among genetic epilepsies, variants in sodium channel coding genes constitute a major subgroup. Variants in *SCN8A*, the coding gene for Na_V_1.6 channels, are characterized by a variety of symptoms including intractable epileptic seizures, psychomotor delay, progressive cognitive decline, and others such as autistic features, ataxia or dystonia. Standard anticonvulsant treatment has only limited impact on the course of disease.

**Experimental Approach:** Personalized therapeutic regimens tailored to disease-causing pathophysiological mechanisms may offer the specificity required to overcome intractability. Toward this aim, we investigated *in vitro* in neuroblastoma cells the effects of S-Licarbazepine, a third-generation dibenzazepine and enhancer of slow inactivation of voltage gated sodium channels, on three gain-of-function Na_V_1.6 variants linked to representative phenotypes of mild epilepsy (G1475R), developmental and epileptic encephalopathy (M1760I) and intellectual disability without epilepsy (A1622D).

**Key Results:** S-Licarbazepine strongly enhances the slow and - less pronounced – the fast inactivation of Na_V_1.6 wildtype channels. It acts similarly on all tested variants and irrespective of their particular biophysical dysfunction mechanism. Beyond that S-Licarbazepine has variant-specific effects including a partial reversal of pathologically slowed fast inactivation dynamics (A1622D, M1760I) and a trend to reduce the enhanced persistent Na^+^ current by A1622D variant channels.

**Conclusion and Implications:** These data bring out that S-Licarbazepine not only owns substance-specific effects, but also holds variant-specific effects, which can variably contribute to functional compensation of distinct channel-specific biophysical properties and thereby highlighting the role of personalized approaches, which likely will be key to improved and successful treatment not only of *SCN8A*-related disease.

**Bullet points:**

- **What is already known?** S-Lic strongly modulates slow and - to a less extend - fast inactivation of wild-type Na_V_1.6 channels.
- **What this study adds?** Differential modulatory effects of S-Lic extend to Na_V_1.6 A1622D, M1760I and G1475R variant channels irrespective of their leading biophysical mode of gain-of-function and variably contribute to variant-specific functional compensation of their altered biophysical properties.
- **Clinical significance:** These data suggest therapeutic potential of S-Lic for *SCN8A* neuropsychiatric disorders and highlight the role of personalized approaches aimed at increasingly precise correction of underlying pathophysiological mechanisms.

## Introduction

New technology such as next generation sequencing has led to the identification of a growing number of associated *de novo* genetic variants, providing the basis for subsequent pathophysiological studies. At the same time, the unfolding complexity of the pathophysiological landscape of genetic epilepsies emerges as one likely reason for limited therapeutic success experienced with standard care.

Sodium channelopathies have initially been identified as one of the most frequent causes of genetic forms of epilepsy. *SCN8A* – the gene encoding the human Na_V_1.6 voltage gated sodium channel (VGSC) – has been recognized as an epilepsy-associated gene in 2012 (Veeramah et al., 2012). Subsequently, many pathogenic *de novo SCN8A* variants have been described in epileptic patients displaying phenotypes ranging from mild epilepsy (e.g. benign familial infantile seizures) to severe developmental and epileptic encephalopathies (DEEs, (Gardella & Moller, 2019; Gertler & Carvill, 2019; Johannesen et al., 2021) including non-epileptic symptoms such as intellectual disability (O’Brien & Meisler, 2013). To cover the full phenotypic and biophysical spectrum we chose two variants with biophysical and neuronal gain-of-function (GOF, M1760I, G1475R) and one with biophysical GOF but neuronal loss-of-function (LOF, A1622D), as outlined above (Fig. 1). While M1760I manifests a severe phenotype with intrauterine onset of seizures, epileptic encephalopathy, intellectual disability, hypotonia and blindness, A1622D is associated with developmental delay, intellectual disability, autism, and motor impairment but not epilepsy (Liu et al., 2019). G1475R has independently been described in several patients (Gardella et al., 2018; Liu et al., 2019; Parrini et al., 2017; Wang et al., 2017; Xiao et al., 2018) presenting with a wide phenotypic spectrum ranging from benign epilepsy to DEE with intellectual disability and motor impairment. Recent work by our group characterizing multiple genetic *SCN8A* variants demonstrated clear genotype-phenotype-correlations based on gain- or loss-of-function of Na_V_1.6 channels and associated neuron dysfunction (Liu et al., 2019): While increased firing always correlated with GOF mechanisms of epilepsy-causing variants (M1760I, G1475R and others), reduced firing in patients without epilepsy only partially accounted for biophysical LOF mechanisms. This became obvious specifically for one variant (A1622D), which was found to decrease neuronal excitability although presenting with a dramatic slowing of fast inactivation of channel kinetics (resembling a GOF mechanism). This effect was so pronounced that neurons failed to properly repolarize and were driven into depolarization block with decreased AP firing, demonstrating the conversion of biophysical channel GOF into LOF on the level of neuron excitability. These data raise the question, whether patients with a “LOF-phenotype” (without epilepsy) could also benefit from treatment with sodium channel blockers (SCBs) and indicate that successful pharmacologic intervention will require drugs designed to specifically address GOF or LOF mechanisms to (largely) restore physiological channel and neuron function.

**Figure 1:**
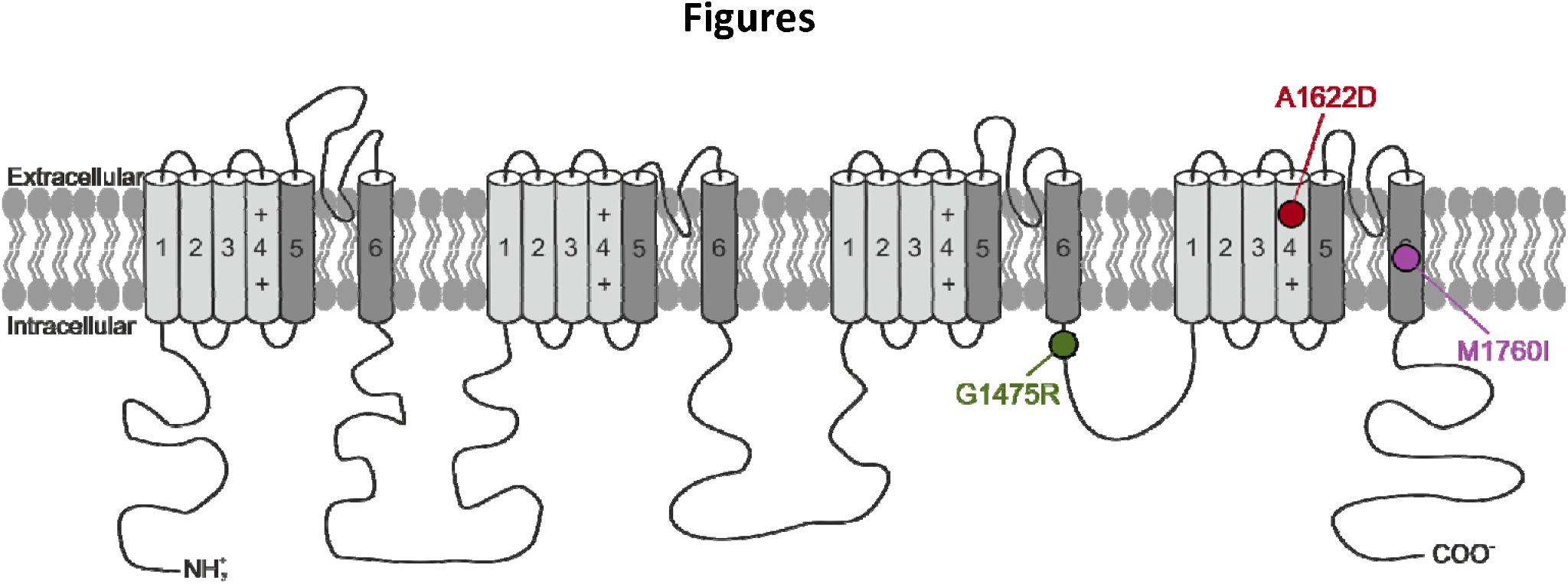
The localization of the three variants in the human Na_V_1.6 channel investigated in this study (M1760I, G1475R, A1622D).

Eslicarbazepine acetate (ESL) is a third-generation member of the dibenzazepine family of antiseizure medication (ASM) and is FDA and EMA approved as adjunctive therapy in adults with focal-onset seizures. As prodrug it undergoes rapid presystemic metabolic hydrolysis to eslicarbazepine (S-Lic), also known as S-Licarbazepine (Bialer & Soares-da-Silva, 2012). Like SCBs within the same pharmacological family (i.e. carbamazepine (CBZ) and oxcarbazepine (OXC)) S-Lic exhibits enhanced binding selectivity to the inactive state of VGSCs, but enhances rather slow inactivation of VGSC than fast inactivation as known for CBZ and OXC (Soares-da-Silva et al., 2015). Importantly, S-Lic maintains significant add-on effects to CBZ, such as antiepileptogenic effects (Doeser et al., 2015), subtype-selectivity to Na_V_1.2 and Na_V_1.6 isoforms (Holtkamp, Opitz, Hebeisen, Soares-da-Silva, & Beck, 2018) and has proven effective and well-tolerated as adjunctive therapy in patients refractory to standard ASM (Shorvon et al., 2017).

Together, these findings suggest that S-Lic may hold great potential for patients with select VGSC gain-of-function variants such as underlying *SCN8A*-related neuropsychiatric disorders.

## Methods

### Experimental protocols and design

#### 1) Mutagenesis

The constructs of the coding sequence (cDNA) of human Na_v_1.6 (WT), A1622D, G1475R, and M1760I α-subunit channel engineered in a Tetrodotoxin (TTX)-resistant channel isoform (so that the current obtained from the transfected channels can be isolated after the addition of TTX), and the constructs of human β_1_- and β_2_-subunits of VGSC (modified to express enhanced green fluorescent protein or a CD8 marker respectively) where acquired and modified as described by (Liu et al., 2019).

#### 2) Transfection and expression in ND7/23 cells

ND7/23 cell line was purchased from Sigma Aldrich. The cells were cultured at 37 °C, with 5 % CO_2_ humidified atmosphere in Dulbecco’s modified Eagle nutrient medium (DMSO, Invitrogen) complemented by the addition of 10 % fetal calf serum (PAN-Biotech) and 1 % L-glutamine 200 mM (Biochrom). One day before the transfection, ND7/23 cells were plated in 35 mm petri dishes so that cells will be 70 - 90 % confluent at the time of transfection. At the day of transfection, 4.4 μg of cDNA (4 μg of the TTX resistant α-subunit ‘WT or mutant human *SCN8A* cDNAs’, and 0.2 μg of each of the human β_1_- and β_2_-subunits of VGSC), along with Lipofectamine 2000 (Invitrogen) were diluted in 2 separate tubes in 250 μl of Opti-MEM I Reduced Serum Medium (Invitrogen), mixed gently and incubated for 5 minutes. After the 5-minute incubation, the diluted

DNA and diluted Lipofectamine 2000 were combined and incubated for 20 minutes at room temperature, then added to the dishes dropwise. Voltage-clamp recordings for the cells expressing all the three subunits were performed 48 hours after transfection. Those cells were recognized by the presence of TTX-resistant Na^+^ current, green fluorescence and anti-CD8 antibody coated microbeads on the cell surface (Dynabeads M450, Dynal) indicating the presence of α-subunit, β_1_-subunit, and β_2_-subunit respectively.

#### 3) Electrophysiology

Standard whole-cell recordings were performed using an Axopatch 200B amplifier, and a DigiData 1320A digitizer. The amplifier head-stage was mounted on a motorized mini manipulator (LN Unit Junior, Luigs & Neumann Feinmechanik und Elektrotechnik GmbH, Germany) and the cells were visualized with an inverted microscope (Axiovert A1, Zeiss). Leakage and capacitive currents were subtracted using a prepulse protocol (−P4). All recordings were performed at room temperature of 21-24 °C.

##### Voltage-clamp recordings

Recordings in transfected ND7/23 cells were performed 10 minutes after establishing the whole-cell configuration. Only cells with a peak Na^+^ current of at least 1 nA and a maximal voltage error due to residual series resistance less than 5 mV after 90 % compensation were chosen for further evaluation. 500 nM TTX + 300μM of S-Lic or vehicle (DMSO) were added to the bath solution to block all endogenous Na^+^ currents. Our *in vitro* studies were conducted with and without administration of 300 µM S-Lic, a concentration that can be warranted considering the high lipid:water partition coefficient of S-Lic (50:1) and its low affinity for non-specific protein binding (Hebeisen et al., 2015). Previous estimates of the concentration of S-Lic at the cerebrospinal fluid of some healthy volunteers resulted in peak values even higher to the concentration used for the *in vitro* experiments in this study (Galiana, Gauthier, & Mattson, 2017). Currents were filtered at 5 kHz and digitized at 20 kHz. Cells were held at −100 mV. Borosilicate glass pipettes with a final tip resistance of 1.5–3 MΩ when filled with internal recording solution. For voltage clamp recordings, the pipette solution contained (in mM): 10 NaCl, 1 EGTA, 10 HEPES, 140 CsF (pH was adjusted to 7.3 with CsOH, osmolarity was adjusted to 310 mOsm/kg with mannitol). The bath solution contained (in mM): 140 NaCl, 3 KCl, 1 MgCl_2_, 1 CaCl_2_, 10 HEPES, 20 TEACl (tetraethylammonium chloride), 5 CsCl and 0.1 CdCl_2_ (pH was adjusted to 7.3 with CsOH, osmolarity was adjusted to 320 mOsm/kg with mannitol).

##### Drugs

Elicarbazepine (S-Lic; BIA 2-194, BIAL - Portela & Ca) was dissolved in DMSO. In all pharmacological experiments, control recordings contained equal amounts of DMSO (0.1 %). All chemicals for solutions were acquired from Sigma-Aldrich, unless stated otherwise. All studies were conducted with 300 μM S-Lic according to previous publications (Doeser, Soares-da-Silva, Beck, & Uebachs, 2014).

### Data acquisition and statistical analysis

The biophysical parameters of human WT and mutant Na_v_1.6 channels were obtained as described previously by (Liu et al., 2019). Detailed descriptions can be found in the supplementary methods section.

Data was recorded using pCLAMP 8 data acquisition software (Molecular Devices) and analysed using Clampfit software of pCLAMP 11.0.3 (Axon Instruments), Microsoft Excel (Microsoft Corporation, Redmond, WA, USA) and GRAFIT 3.01 software (Erithacus UK). Statistics were performed using Graphpad software V7 (Graphpad prism, San Diego, CA, USA). All data was tested for normal distribution using Shapiro–Wilk test. For comparison between two groups (+/-S-Lic), unpaired *t-*test was used for normally distributed data and Mann-Whitney U-test was used for non-normally distributed data. For comparison between multiple groups (WT+/-S-Lic and variant +/-S-Lic) one-way ANOVA with Dunnett’s *post-hoc* test was used for normally distributed data and ANOVA on ranks with Dunn’
ss *post-hoc* test for non-normally distributed data. All data are shown as mean ± standard error of the mean (SEM), n indicates the number of cells. Statistical significance requires the p value to be less than 0.05. For all statistical tests, significance is indicated on the figures using the following symbols: * p < 0.05, ** p < 0.01, *** p < 0.001, **** p < 0.0001.

## Results

### S-Lic strongly modulates slow inactivation of wild-type Na_V_1.6 channels

To confirm previously reported effects of S-Lic (Hebeisen et al., 2015; Holtkamp et al., 2018) in our preparations we performed whole-cell patch-clamp recordings in ND7/23 neuroblastoma cells transfected with the TTX-resistant Na_V_1.6 wild-type α-subunit. Recordings were performed in presence of TTX, cesium, TEA and cadmium to block endogenous voltage gated sodium-, potassium- and calcium channels. In line with previous publications S-Lic predominately modulated the voltage dependence of steady-state slow inactivation (Vehicle: V_1/2_ = −55.96 ±1.16 mV, n=7; Vehicle + 300 μM S-Lic: V_1/2_ = −72.06 ±0.71 mV, n=7; p<0.0001) by substantially shifting V_1/2_ toward hyperpolarized potentials and thereby reducing the area under the curve (AUC) considerably (Fig. 2G). All other investigated biophysical parameters were either affected markedly less pronounced (hyperpolarizing shift of the steady-state fast inactivation, Fig. 2D; accelerated entry into slow inactivation, Fig. 2F; decreased slope of the steady-state slow inactivation, Fig. 2G), showed only a trend (transient and persistent peak currents, Fig. 2B and C; accelerated time course of fast inactivation, Fig. 2E inset), or remained unchanged (voltage dependence of steady-state activation, Fig. 2D; recovery from fast inactivation, Fig. 2E).

**Figure 2:**
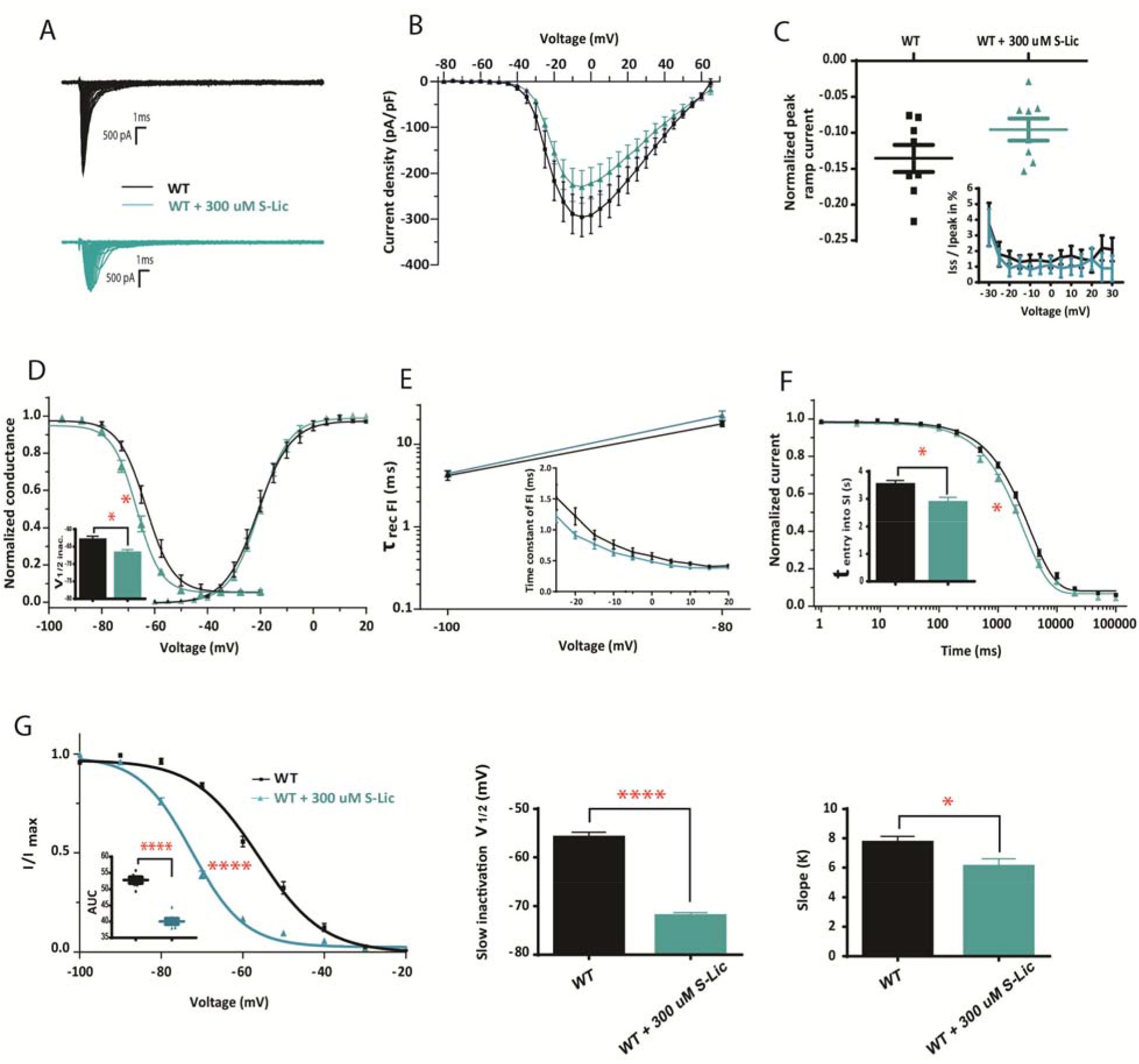
The major effects of Eslicarbazepine (S-Lic) on Na_v_1.6 wild-type channels. WT channels were transfected into ND7/23 cells for voltage clamp recordings. Na^+^ currents recorded in the presence of 300 μM of S-Lic (turquoise) or vehicle (black), accompanied by tetrodotoxine (TTX) in order to block the endogenous Na^+^ channels. **(A)** Representative Na^+^ current traces from WT (black) and WT + 300 μMS-Lic (turquoise). **(B)** Peak transient Na^+^ currents normalized by cell capacitances plotted against voltage. WT, n=14. WT + 300 μM S-Lic, n=9. **(C)** Ramp current peaks (persistent current) normalized to transient peak currents I_peak_ upon ramp stimuli from −100 to +40 mV lasting 800 ms, n=8. Inset of figure: Persistent Na^+^ currents (I_ss_, for the “steady-state” current) normalized to the transient peak current (I_ss_/I_peak_) plotted against voltage, n=12 for WT, and n=8 for WT + 300 μM S-Lic. **(D)** Voltage-dependent steady state activation and inactivation curves in WT Na_v_1.6 in the presence of 300 μM of S-Lic or vehicle, lines represent Boltzmann functions fit to the data points. Exposure to 300 μM of S-Lic does not alter the voltage dependence of activation but causes a hyperpolarizing shift (vehicle, V_1/2_ = −63.21 ±1.2 mV, 300 μM S-Lic, V_1/2_ = −66.98 ± 1.08 mV) in the voltage dependence of fast inactivation. Activation: n=10 for WT, and n=9 for WT+ 300 μM S-Lic. Inactivation: n=10 for WT, and n=7 for WT + 300 μM S-Lic. **(E)** Mean values of the recovery time constant at two different holding voltages (−80 and −100 mV, n=9 for WT, and n=11 for WT + 300 μM S-Lic). Inset of figure: Voltage-dependence of the major time constant of fast inactivation (n=10 for WT, and n=9 for WT + 300 μM S-Lic). **(F)** Entry into slow inactivation. The lines represent fits of a first-order exponential function for the data points. Exposure to 300 μM of S-Lic accelerates the entry into slow inactivation (τ_entry_ = 2847.06 ±208.4 ms, n =11) when compared to vehicle (τ_entry_ = 3500.26 ±170.8 ms, n=10). **(G)** Steady-state slow inactivation. A standard Boltzmann function was fit to the data points. Exposure to 300μM of S-Lic causes a hyperpolarizing shift of the steady-state slow inactivation (n=7 for Vehicle, V_1/2_ = −55.96 ±1.16 mV and n=7 for 300 μM S-Lic, V_1/2_ = - 72.06 ±0.71 mV), and decreased the slope of the steady-state slow inactivation when compared to vehicle (vehicle, k = 7.69 ±0.44, 300 μM S-Lic, k = 6.05 ±0.54, n=7). Inset of figure: The area under the curve was determined and respective values were significantly lower for WT + 300 μM S-Lic group (vehicle, AUC = 52.8 ±0.83, 300 μM S-Lic, AUC = 40.05 ± 0.84, n=7). Shown are means ± SEM for each data point. n, number of recorded cells, * p <0.05, **** p < 0.0001 (unpaired *t*-test or Mann-Whitney U-test).

### S-Lic modulation of slow inactivation extends to Na_V_1.6 A1622D, M1760I and G1475R variant channels irrespective of their leading biophysical mode of GOF

To investigate whether S-Lic conveys similar modulatory effects on Na_V_1.6 variants analog to what has been demonstrated for slow inactivation gating properties and kinetics of wild-type channels we next performed voltage-clamp experiments in ND7/23 cells transfected with M1760I, G1475R or A1622D α-subunits. In line with previous reports by our group (Liu et al., 2019), baseline recordings confirmed a hyperpolarizing shift of the activation curve (M1760I, Suppl. Fig. 1A), a depolarizing shift of the steady state fast inactivation curve (G1475R, Suppl. Fig. 1E) and a pronounced slowing of fast inactivation kinetics (A1622D, Fig. 3C) with large ramp currents and correspondingly increased persistent Na^+^ currents as the main biophysical features (Fig. 3R) of the respective variants.

**Figure 3:**
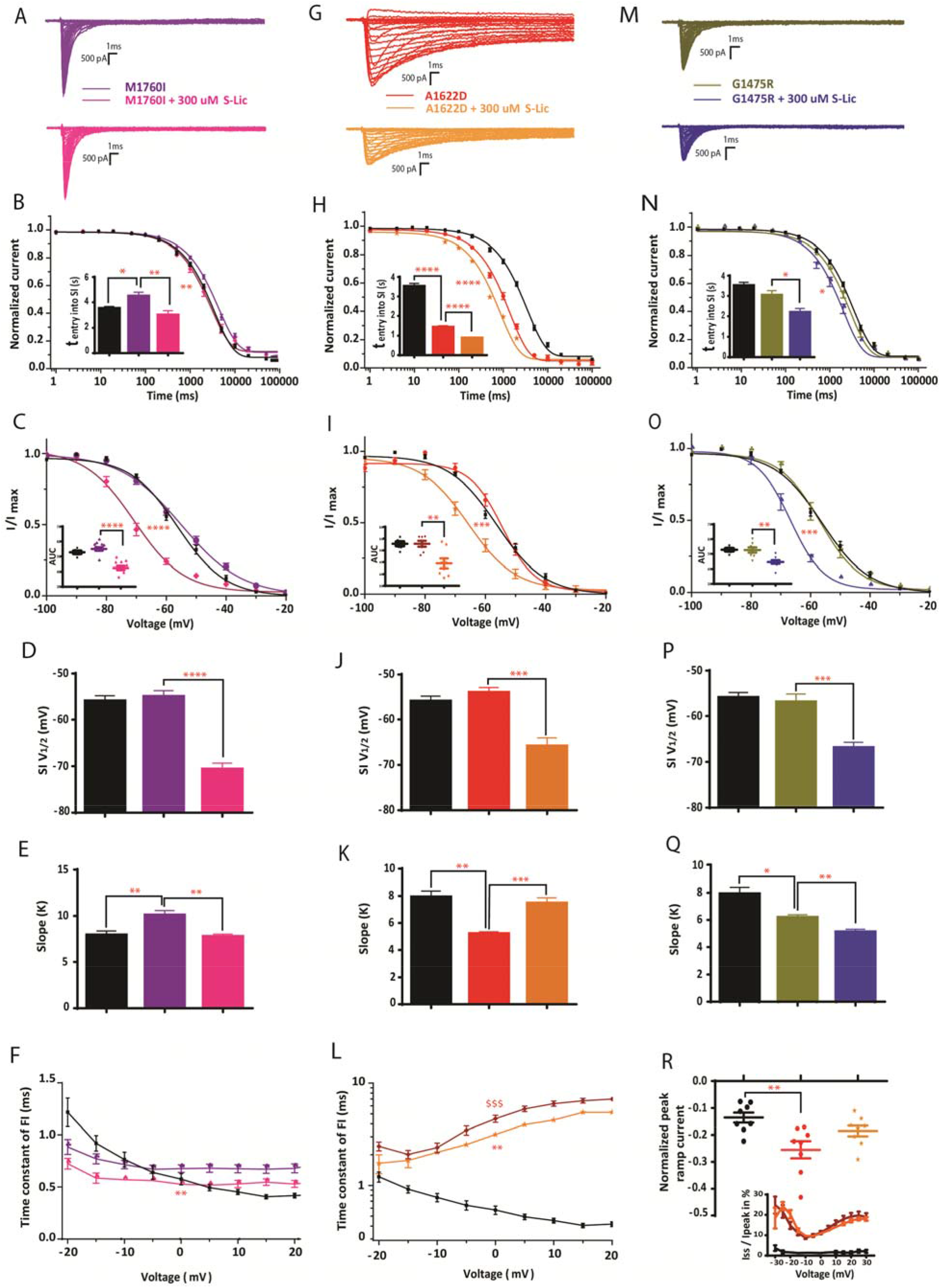
Effects of Eslicarbazepine (S-Lic) on various Na_v_1.6 mutant channels expressed in the rodent neuroblastoma cell line ND7/23. Na^+^ currents were recorded in the presence of 300 μM of S-Lic or vehicle, accompanied by tetrodotoxine (TTX) in order to block the endogenous Na^+^ channels. **(A)** Representative Na^+^ current traces from M1760I (violet), and M1760I + 300 μM S-Lic (pink). **(B)** Entry into slow inactivation in variant M1760I channels. Exposure to 300 μM of S-Lic in M1760I has accelerated the entry into slow inactivation (τ_entry_ = 2984.18 ±355.44 ms, n=10) when compared to vehicle (τ_entry_ = 4474.82 ±322.82 ms, n=12). **(C-E)** Steady-state slow inactivation in variant M1760I channels. Inset of figure: The area under the curve (AUC) was determined and respective values were significantly lower for M1760I + 300μMS-Lic group (vehicle, AUC = 55.01 ±1.15, 300μMS-Lic, AUC = 42.46 ±1.36). Exposure to 300 μM of S-Lic in M1760I has caused a hyperpolarizing shift of the steady-state slow inactivation (V_1/2_ = −70.66 ±1.35 mV), when compared to vehicle (V_1/2_ = −54.98 ±1.28 mV) (D), and a decrease in the slope of the steady-state slow inactivation (M1760I +300μMS-Lic; k = 7.7 ±0.72; M1760I; k = 10.07 ±0.52) (E) M1760I n=10, M1760I + 300μMS-Lic n=9. **(F)** Voltage-dependence of the major time constant of fast inactivation in variant M1760I channels. S-Lic significantly reversed M1760I variant-specific mild slowing of fast inactivation, as reflected by a partial normalization of the otherwise increased time constant of fast inactivation at 0 mV (M1760I, τ at 0 mV = 0.67 ± 0.03ms, n= 10, and M1760I + 300 µM S-Lic, τ at 0 mV = 0.52 ±0.02 ms, n=10). **(G)** Representative Na^+^ current traces from A1622D (red), and A1622D + 300 μM of S-Lic (orange). **(H)** Entry into slow inactivation in variant A1622D channels. Exposure to 300 μM of S-Lic further accelerates the entry into slow inactivation vs vehicle (n=8 for A1622D τ_entry_ = 1422.56 ±93.3ms, n=10 for A1622D + 300μM S-Lic τ_entry_ = 871.1 ±53.5 ms). **(I-K)** Steady-state slow inactivation in variant A1622D channels. Inset of figure: The area under the curve was determined and respective values were significantly lower for A1622D + 300 μM S-Lic group (vehicle, AUC = 52.8 ±1.27, 300 μM S-Lic, AUC = 44.5 ±2.06). Exposure to 300 μM of S-Lic causes a hyperpolarizing shift of the steady-state slow inactivation (A1622D, V_1/2_ = −54.0 ± 1.11 mV, A1622D + 300μMS-Lic, V_1/2_ = −65.83 ±1.8 mV) (J), and a decrease in the slope of the steady-state slow inactivation (A1622D, k = 5.2 ± 0.15, A1622D + 300 μM S-Lic, k =7.46 ±0.4) (K). A1622D n=7, A1622D + 300 μM S-Lic n=9 **(L)** Voltage-dependence of the major time constant of fast inactivation in variant A1622D channels. Both S-Lic and without S-Lic A1622D groups show a little voltage dependence versus WT, however, the +S-Lic group shows a significant acceleration in the transition from the activated to the fast inactivated state in comparison to without S-Lic group (n=9 for A1622D; τ at 0 mV = 4.49 ±0.35 ms, and n=9 for A1622D + 300μMS-Lic; τ at 0 mV = 3.15 ±0.20 ms). **(M)** Representative Na^+^ current traces from G1475R (olive green), G1475R + 300μM of S-Lic (blue). **(N)** Entry into slow inactivation in variant G1475R channels. Exposure to 300 μM of S-Lic accelerates the entry into slow inactivation (n=13 for G1475R τ_entry_ = 3028.89 ±234.29 ms, and n=13 for G1475R + 300μM S-Lic τ_entry_ = 2182.25 ±202.39 ms). **(O-Q)** Steady-state slow inactivation in variant G1475R channels. Inset of figure: The area under the curve was determined and respective values were significantly lower for G1475R + 300 μM S-Lic group (vehicle, AUC = 52.63 ±1.73, 300 μM S-Lic, AUC = 44.52 ±1.22). Exposure to 300 μM of S-Lic caused a hyperpolarizing shift of the steady-state slow inactivation (G1475R, V_1/2_ = −56.93 ±1.8 mV, and for G1475R + 300 μM S-Lic, V_1/2_ = −66.93 ±1.27 mV) (P), and further decreased the slope of the steady-state slow inactivation (G1475R k = 6.17 ±0.20, and G1475R + 300 μM S-Lic, k = 5.11 ±0.2) (Q). G1475R n=8, G1475R + 300 μM S-Lic n=9. **(R)** Ramp current peaks (persistent current) in variant A1622D channels normalized to transient peak currents I_peak_. There was a large ramp current for the A1622D variant corresponding to a high persistent current and exposure to 300 μM of S-Lic had shown a trend to lower it # p = 0.0835 (A1622D = - 0.25 ±0.03, n= 8, A1622D + 300 μM S-Lic = −0.18 ±0.01, n=8). Inset of figure: Persistent Na^+^ currents (I_ss_, for the “steady-state” current) normalized to the transient peak current (I_ss_/I_peak_) plotted against voltage, n=9 for A1622D and A1622D + 300 μM S-Lic. Shown are means ± SEM for each data point. WT (black), M1760I (violet), M1760I+300 μM S-Lic (pink), A1622D (red), A1622D + 300 μM of S-Lic (orange), G1475R (olive green), and G1475R + 300 μM of S-Lic (blue). Shown are means ±SEM for each data point. n, number of recorded cells; * p < 0.05; ** p < 0.01; *** p < 0.001; **** p < 0.0001; the same goes for $ sign but indicates a difference vs. WT. Unpaired t-test or Mann-Whitney U-test were used for +/-S-Lic comparisons, for multiple group comparisons (WT+/-S-Lic and variant+/-S-Lic) one-way ANOVA with Dunnett’s *posthoc* test or ANOVA on ranks with Dunn’
ss *posthoc* test were used.

300 μM S-Lic proved active on all three tested variant channels, regardless of their heterogeneous functional characteristics. As observed for the wild type, S-Lic significantly shifted steady-state slow inactivation curves about 10 to 15 mV to more hyperpolarized potentials (M1760I: almost −15 mV, Fig. 3C+D; G1475R: −10 mV, Fig. 3O+P; A1622D: almost −12 mV, Fig.3I+J and Suppl. Table 1) and reduced the slope factor k of steady-state slow inactivation in a comparable manner as seen for the wildtype (Fig. 2G, Δk = −1.6) except for the A1622D variant channels (M1760I: Δk = −2.3, G1475R: Δk = −1.1, A1622D: Δk = +2.3, Fig.3E, Q, K and Suppl. Table 1). Furthermore, S-Lic accelerated the entry into slow inactivation for all variants to a variable extent between approximately 28 % (G1475R) and 33 % (M1760I) up to 39 % (A1622D), which outlines a much stronger effect on variant channels than seen on wildtype channels (19 %; Suppl. Table 2). Moreover, the strongest effect in absolute time units – an earlier entry into the slow inactivation of almost 1500 ms – was seen in M1760I variant channels which had compared to the wildtype a previously significantly prolonged entry into slow inactivation (WT: 3500 ms ±170.8 vs. M1760I: 4475 ms ±322.5, Suppl. Table 2) demonstrating that S-Lic can more than compensate the variant effect for this biophysical feature.

The described modulation of slow inactivation by S-Lic is generally considered the main mechanism underlying its anticonvulsant properties via promoting the use-dependent transition of the activated to the inactivated state, even though impaired slow inactivation may not actually be part of the altered biophysical properties of a channel variant. As already demonstrated by Liu et al., 2019 slow inactivation curves indeed were not shifted for any of the three variants, investigated in this work. Instead, each variant associated with a combination of multiple other biophysical changes (Liu et al., 2019). Whether S-Lic may exhibit additional effects on variant channels that are distinct from the ones observed on wildtype Na_V_1.6 may critically impact its suitability for the treatment of affected patients.

### Differential modulatory effects of S-Lic variably contribute to compensation of functionally altered Na_V_1.6 M1760I, G1475R and A1622D channels

To explore this possibility, we next asked whether S-Lic may have additional and Na_V_1.6 variant-specific effects, which may directly interject and partially reverse or, alternatively, aggravate certain aspects of dysfunctional M1760I, G1475R and A1622D channels. To address these questions a broad spectrum of biophysical parameters was investigated before and after application of 300 μM S-Lic, followed by subsequent comparative analysis. These data were put in direct context with the respective values obtained for wild type channels in absence of S-Lic. Recordings were performed in ND7/23 cells as described above.

#### A1622D

While transient peak Na^+^ currents were insensitive to S-Lic and comparable among the three variants and the wild type (Fig. 2A+B, Fig. 3A+G+M, Suppl. Fig. 1H, Suppl. Table 1), S-Lic caused a trend to reduced persistent Na^+^ currents of A1622D channels. This effect was specific to the A1622D variant and was therefore (likely) independent of typical S-Lic mediated modulation of the voltage-dependence of slow inactivation (normalized ramp current peaks: WT = −0.13 ±0.01, A1622D = −0.25 ±0.03, A1622D + 300 µM S-Lic = −0.18 ±0.01, p = 0.08; Fig. 3R). As expected from wildtype experiments, our data revealed that S-Lic neither interferes with voltage dependence of activation of M1760I channels nor G1475R channels, but demonstrated a significant variant-specific shift of steady-state activation of A1622D channels (A1622D + 300 µM S-Lic = −6.60 ±0.95 vs. A1622D = −10.07 ±1.11, p = 0.03). Notably, unlike observed for wildtype channels, S-Lic may convey additional effects resulting in an adverse amplification of certain GOF aspects of variant channels, as demonstrated for A1622D by an add-on increase of the already preexisting depolarized shift of the steady-state fast inactivation curve (A1622D V_1/2_ = −56.31 ±1.41 mV, A1622D + 300 μM S-Lic V_1/2_ = −51.85 ±0.6 mV, p = 0.005, Suppl. Fig. 1C, Suppl. Table 2).

#### M1760I

As already reported earlier on in this study, M1760I predominantly causes a hyperpolarizing shift of the activation curve (Suppl. Fig. 1A) and a slowing of the fast inactivation kinetics (Suppl. Fig 1F). Whereas no effect of S-Lic on steady-state activation was seen, S-Lic significantly reversed M1760I variant-specific mild slowing of fast inactivation, as reflected by a partial normalization of the otherwise increased time constant of fast inactivation at 0 mV (WT: τ at 0 mV = 0.57 ± 0.05 ms, M1760I: τ at 0 mV = 0.67 ± 0.03ms, M1760I + 300 µM S-Lic: τ at 0 mV = 0.52 ±0.02 ms; Fig. 3F and Suppl. Table 2).

While a slightly earlier entry into slow inactivation of M1760I variant channels could be rescued by S-Lic – we did not find any clear indication of impaired slow inactivation for any of the other variants (neither in this study nor in the preceding work by Liu et al., 2019). We nevertheless noticed subtle changes of the slopes of steady-state slow inactivation of the variants in comparison to the WT, which variably responded to S-Lic. While for A1622D and M1760I the slopes of the steady-state slow inactivation indeed reverted to WT levels (Fig. 3E+K), similarly to the observation with wild type channels the slope slightly further decreased for G1475R (Fig. 3Q). However, these small modulatory effects are present in addition to the major S-Lic induced hyperpolarizing shifts of V_1/2_ of steady-state slow inactivation described above and seem to be part of a complex loss-of-function effect comprising a highly significant decreased AUC (p < 0.0001) of the steady-state slow inactivation curve and therefore as individual effect of minor relevance. S-Lic did not alter the recovery from fast inactivation at −80mV holding potential but an acceleration was observed at −100mV (at −100mV: WT τ_Rec_ = 4.16 ±0.52; M1760I τ_Rec_ = 3.95 ±0.35, M1760I + 300 µM S-Lic τ_Rec_ = 2.79 ±0.14, p = 0.049, Suppl. Fig.1B).

*G1475R*: As seen for the wildtype and much stronger for the two other variants the G1475R channels showed also a trend to faster inactivation kinetics (p = 0.68) with a small reduction of the time constant of fast inactivation at 0 mV comparable to the wildtype (Suppl. Fig. 1F, Suppl. Table 2). Apart from that, S-Lic did not reverse the variant-specific shift of the steady-state fast inactivation curve, although a significant shift was seen for wildtype channels (Suppl. Fig. 1E, Suppl. Table 1).

Taken together, these data indicate additional variant-specific mechanisms of S-Lic, some of which are likely contributing to overall anticonvulsant properties while others may rather mediate moderate counterbalancing or converse effects in the context of certain mutant channels.

A comprehensive summary of the biophysical properties of wild type and variant channels and respective modulatory effects of S-Lic investigated in this study is provided in supplementary tables 1 and 2.

## Discussion

Our biophysical findings demonstrate the previously described enhancement of the slow and fast inactivation (SI and FI) comprising an earlier entry into SI and a strong hyperpolarizing shift of the SI curve in WT Na_v_1.6 channels, which could be similarly observed for all variant channels. These findings indicate that none of the variants interfered with the S-Lic binding site and give no hints to “on-target-resistances”. Therefore, we see the observed effects independent of the distinct variant effects and of the severity of the disease.

Whereas the effects of S-Lic are well-known and several studies describing the effects of S-Lic in heterologous expression systems (Hebeisen et al., 2015) and rat neurons (Holtkamp et al., 2018) have been performed on wildtype cells, to our best knowledge, systematic data about variant-specific effects of S-Lic are not available and presented with this study for the first time.

Each variant was characterized by a distinct clinical phenotype including severe DEE (M1760I), mild epilepsy (G1475R) or intellectual disability without epileptic seizures (A1622D) – altogether providing a solid representation of the wide phenotypic spectrum of *SCN8A* related disorders – but also by distinct kinetic alterations, predominantly by a hyperpolarizing shift of the activation curve (M1760I), a depolarizing shift of the steady-state fast inactivation curve (G1475R) and a pronounced slowing of fast inactivation kinetics (A1622D) with large ramp currents and correspondingly increased persistent Na^+^ currents. The effects of the latter variant were of particular interest since a neuronal (and clinical) LOF with broadened AP half-width, impaired AP repolarization and consecutively impaired AP firing revealed a dramatic GOF mechanism at biophysical level, leading to a depolarization block on neuronal level and explained in this way the clinical LOF phenotype nicely. Among all variants and the already described substance-specific effects, S-Lic showed the most prominent variant-specific effects on these A1622D channels by a partial reversal of the pathologically slowed FI dynamics and caused a trend to reduce the massively enhanced persistent Na^+^ currents. Based on these results, we speculate that S-Lic could be beneficial for patients carrying this mutation. On the other hand, dissolving of the depolarization block could also lead to paradoxical effects with subsequent seizures, although this never occurred in our patient yet. In the two other variants S-Lic did not impact the steady-state activation but – similar to the A1622D channels - lead to a partial normalization of altered FI kinetics for M1760I channels. Additionally, few examples for S-Lic mediated small GOF effects were found (e.g. acceleration of recovery from fast inactivation at −100 mV, but not at −80mV for M1760I channels or a small depolarizing shift of steady-state activation curve for A1622D channels), however, considering the strong variant independent S-Lic mechanisms small add-on GOF effects seem to be of limited relevance. A previously described increased persistent current of G1475R channels in HEK293 cells (Zaman, Abou Tayoun, & Goldberg, 2019) could not be observed in our studies, which might be due to different heterologous expression systems.

In the light of these biophysical data our study would benefit from a clinical correlation with drug history and actual treatment outcome of patients carrying the respective genetic variants. Yet, such an analysis is hampered by the *de novo* occurrence of the variants with only very few affected patients (mostly only single individuals) and the retrospective nature of this study. In fact, none of the mutation carriers (already previously clinically described by Liu et al., 2019) had been treated at any point with S-Lic, nor are descriptions available in the literature making a direct retrospective validation impossible. Nevertheless, the robust *in vitro* effect of S-Lic on the M1760I variant would suggest some effectiveness of related sodium channel blockers (e.g. OXC) or others such as phenytoin or lacosamide. In fact, (Zaman et al., 2019) evaluated in a single tertiary center seven patients with different *SCN8A* variants and found that five of seven patients benefitted clinically from SCB, namely phenytoin, oxcarbazepine, lamotrigine and lacosamide. However, almost all responders to one SCB have shown a non-response to another SCB that was given before in the patient’s history pointing out that variant-specific mechanisms play a major role in anticonvulsive treatment underlining the need for targeted treatment approaches. Until now, only a single patient has been reported with the M1760I variant who, in addition, had been diagnosed with an exceptionally severe developmental phenotype with intrauterine onset of seizures. A variety of ASM were tested at the time, and failed to exert sustained anticonvulsive effects, including phenobarbital, levetiracetam, vigabatrine & clobazam (both revealed positive effects on myoclonias), sultiam, the SCB lacosamide (no benefit) and phenytoin for which at least an initial – although not persistent – response on seizures was described. This course of disease certainly represents an extreme case while no precise information regarding duration of treatment, drug dosages, or serum levels of anticonvulsant compounds is available. In contrast, G1475R was reported to be associated with a relatively mild epilepsy phenotype. The patient was initially started on levetiracetam and topiramate; later on, clobazam was added intermittently for treatment of increased seizure frequency but was discontinued again due to noticed concurrent developmental regression. Finally, oxcarbazepine was prescribed and proofed beneficial. Similar effects for the same variant with a good response to oxcarbazepine and lacosamide were also observed by others (Zaman et al., 2019). In contrast, the patient carrying the A1622D variant did not present any seizures and hence no treatment attempt with ASM has been reported.

While a subset of DEE patients with GOF VGSC variants have shown quite good therapeutic responses to standard SCB, in particular to high-dose phenytoin and carbamazepine (Boerma et al., 2016; Dilena et al., 2017; Gardella & Moller, 2019; Moller & Johannesen, 2016; Zaman et al., 2019) pronounced and often dose-dependent side effects (such as cardiac arrhythmias, irreversible cerebellar atrophy, polyneuropathy, fatigue, cognitive impairment; (Ijff & Aldenkamp, 2013)) or insufficient seizure reduction continue to pose frequent and well known therapy limitations. One of the likely reasons for unfavorable effect/side effect profiles is rooted in the rather non-specific modulation of a broad spectrum of VGSC subtypes by classic SCB (Qiao, Sun, Clare, Werkman, & Wadman, 2014). ESL, as the only currently FDA/EMA approved ASM offering increased subtype selectivity (Holtkamp et al., 2018) and antiepileptogenic effects from preclinical data (Doeser et al., 2014) can serve as a prototypical example and proof-of-concept compound for a new generation of yet to be developed target-specific therapeutic agents.

## Conclusions

Extending on previous studies having demonstrated Na_V_1.2 and Na_V_1.6 specific modulation of slow inactivation by S-Lic (Holtkamp et al., 2018) we now investigated S-Lic in context of a set of Na_V_1.6 channel variants associated with *SCN8A*-related neuropsychiatric disorders. Intriguingly, our experiments identified additional variant-specific mechanisms, distinct from known generic effects on slow inactivation. In summary, these data suggest therapeutic potential of S-Lic for *SCN8A*-related disease and highlight the role of personalized approaches aimed at increasingly precise correction of pathophysiological mechanisms underlying DEE and other neurodevelopmental disorders.

## Acknowledgements

We thank Bial for the supply of S-Lic and the support of this investigator-initiated trial (IIT) and we thank the European Union for enabling this Erasmus exchange program.

## Data and materials availability

All data associated with this study are presented within this manuscript or in the attached Supplementary Materials.

## Author contribution statement

E.B., S.L., U.H. Y.S. and H.L. designed the study. E.B., Y.L., S.L. and T.W. performed, and analyzed the experiments. E.B., S.L., U.H. and T.W. wrote the manuscript and created the graphs, all authors revised the manuscript for intellectual content and approved the final version.

## Ethics approval statement

All recordings and experiments were done according to regulations of local authorities.

## Permission to reproduce material from other sources

All materials are created newly for this publication, if not otherwise stated.

