## Supplemental Fig.1 and Table 1&2 for "*In vitro* effects of S-Licarbazepine as a potential precision therapy on *SCN8A* variants causing neuropsychiatric disorders"

**Supplementary methods**

**Voltage clamp protocols**

The activation curve (conductance versus voltage) was acquired by determining the peak Na^+^ current at various step depolarizations of the measured cells starting from the holding potential of -100 mV. The data obtained was fitted by a Boltzmann function: g/g_max_ (V) = 1/(1+exp[(*V*-*V*_1/2_)/k_V_]), where the conductance (g) is calculated from g = I/(V-V_rev_), g_max_ the maximal conductance, V_rev_ the reversal potential of Na^+^, V_1/2_ the voltage of half-maximal activation, and k_V_ is a slope factor. Steady-state inactivation was determined by measuring the peak current obtained using 100 ms conditioning pulses to various potentials (starting from -110 mV up to -20 mV) followed by the test pulse to -10 mV. The peak current obtained in this protocol reflects the percentage of non-inactivated channels. Then, the data obtained was fitted by a standard Boltzmann function: I/I_max_ (V) = 1/(1+exp[(V-V_1/2_)/k_V_]), where the maximal current amplitude is I_max_, V_1/2_ the voltage of half-maximal inactivation, and k_V_ as a slope factor. Kinetics of fast inactivation was evaluated using the data acquired from the protocol used to determine the voltage-dependence of activation, which was also used to determine the time constants of fast inactivation. To fit to the time course of fast inactivation during the first 70 ms after onset of the depolarization, a second-order exponential function was used resulting in two time constants. Only the fast time constant, named τ, was considered for analysis because the weight of the second slower time constant was rather small (0-25%). For assessing the recovery from fast inactivation, cells were held at -100mV, then depolarized to -20 mV for 100 ms (in order to inactivate all VGSCs). The cells were then repolarized to either -80 mV or -100 mV for increasing durations followed by a second depolarizing step to -20 mV for 5 ms. The fraction recovered "obtained by dividing the current amplitude by the most negative current amplitude obtained during the test pulse" was plotted against the different holding durations back at -80 or -100 mV. To fit the time course of recovery from inactivation, a first-order exponential function with an initial delay was used yielding the time constant τ_rec_ for the variants tested with the exception of the A1622D variant, since a second-order exponential function with an initial delay was considered more fitting (τ_rec fast_). Cumulative protocols were used to assess the entry into, and steady-state slow inactivation (Alekov, Peter, Mitrovic, Lehmann-Horn, & Lerche, 2001). For steady-state slow inactivation, 30 s conditioning pulses starting at –110 mV and stepping by 10 mV gradually up to 0 mV were used, followed by a 100 ms-hyperpolarization to –100 mV (to allow Na^+^ channels to recover from fast inactivation) and then by a 5 ms test pulse at -25 mV. The data was fitted to the Boltzmann function previously mentioned above for fast inactivation. To assess the entry into slow inactivation, the cells were depolarized from the holding potential of –100 mV, to 0 mV for increasing durations, then repolarized again to –100 mV for 100 ms in order to permit the channels to recover from fast inactivation. Then, the cells were depolarized again to 0 mV briefly for 3 ms in order to determine the fraction of slow inactivated channels. Current amplitudes recorded during the test pulses were normalized to the most negative peak obtained and plotted against the initial holding durations. The fitting of the time course of entry into slow inactivation was done using a first order exponential function. The data acquired from the protocol used to determine the voltage-dependence of activation was also used to determine persistent Na^+^ currents. Persistent Na^+^ currents (I_SS_, for the “steady-state‟ current) were determined at the end of the 100 ms depolarizing pulses to different test potentials and normalized to the respective peak current (I_PEAK_) in each sweep. An additional approach was used to visualize the persistent Na^+^ current, a slowly depolarizing ramp stimulus from -100 to +40 mV lasting 800 ms was applied and the data acquired was normalized to the maximum transient peak Na^+^ currents.

**Supplementary Results**

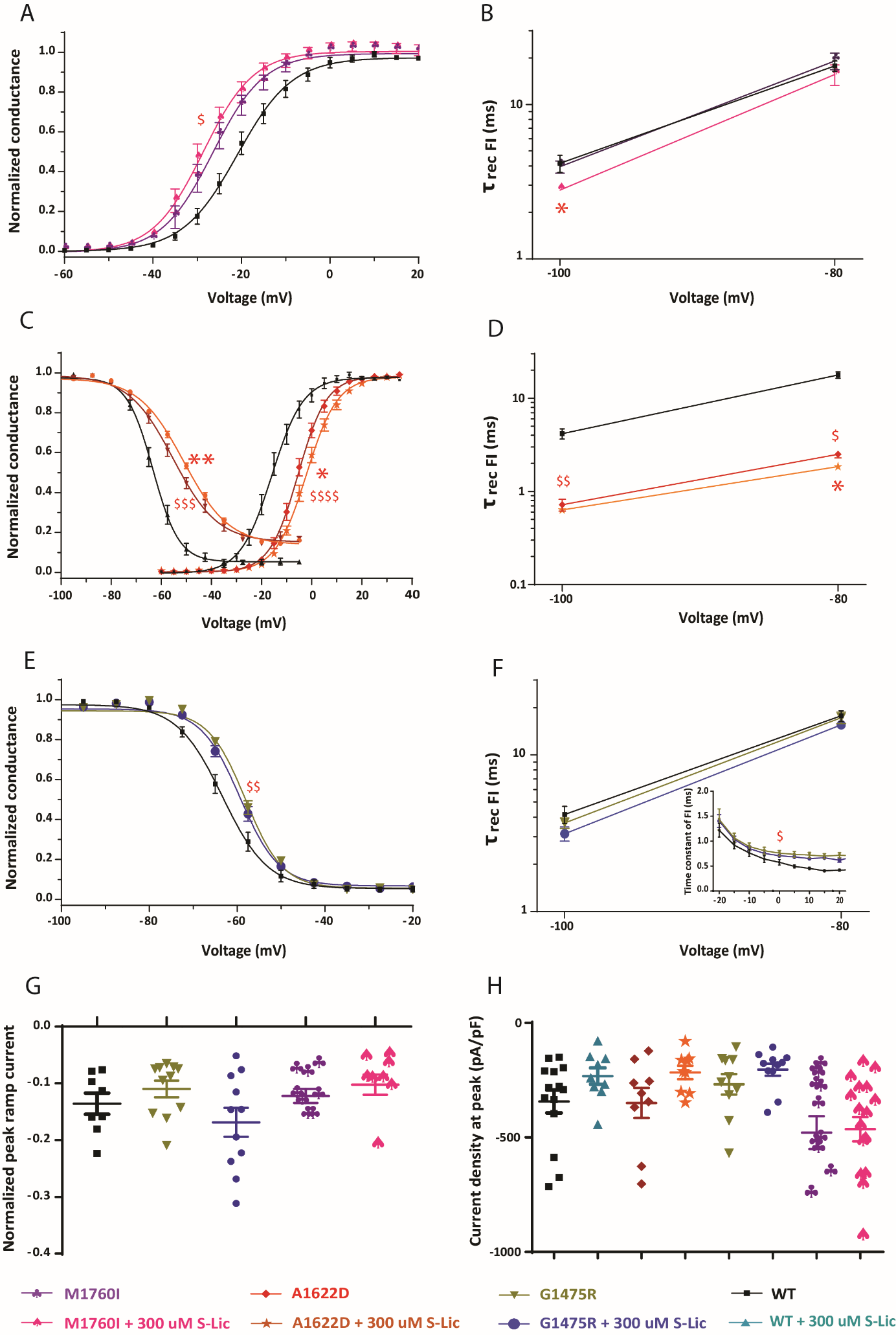

**Supplementary Figure 1: Effects of Eslicarbazepine (S-Lic) on selected biophysical properties of various Na_v_1.6 mutant channels and WT expressed in the rodent neuroblastoma cell line ND7/23. (A)** Steady-state activation curves in M1760I variant Na^+^ channels in the presence of 300 μM of S-Lic (pink) or vehicle (violet). M1760I variant causes a hyperpolarizing shift of the activation curve in M1760I variant channels with respect to the WT and exposure to 300 μM of S-Lic did not seem to alter the voltage-dependence of activation and fast inactivation in M1760I variant. Lines represent Boltzmann functions fit to the data points. **(B)** Mean values of the recovery time constant at two different holding voltages (-80 and -100 mV). S-Lic did not alter the recovery from fast inactivation at -80mV holding potential but an acceleration was observed at -100mV (at -100mV; M1760I τ = 3.95 ±0.35, M1760I + 300 µM S-Lic τ = 2.79 ±0.14, n=7. **(C)** Voltage-dependent steady-state activation and fast inactivation curves of A1622D variant channels. Lines represent Boltzmann functions fit to the data points. Recordings revealed that variant A1622D channels show a depolarizing shift of the activation curve and steady-state fast inactivation curve was also shifted towards depolarized potentials for A1622D channels and the slope was increased compared to WT channels potentials. Exposure to 300μM of S-Lic has shifted both the steady-state activation and fast inactivation curves towards depolarized potentials (Activation: A1622D V_1/2_ = -10.07 ±1.11 mV, n=9, A1622D + 300uM S-Lic V_1/2_ = -6.60 ±0.95 mV, n=9. Fast inactivation: A1622D, V_1/2_ = -56.31 ±1.41 mV, n=7; A1622D + 300 μM S-Lic, V_1/2_ = -51.85 ±0.6 mV, n=10). **(D)** Mean values of the recovery time constant recovery from fast inactivation at two different holding voltages (-80 and -100 mV). A1622D variant channels had an accelerated recovery from fast inactivation at both -80 and -100 mV holding potentials. Exposure to 300 μM of S-Lic had significantly accelerated the time constant for recovery from fast inactivation when compared to vehicle at -80 mV (A1622D τ = 2.5 ±0.21, n=9; A1622D + 300μM S-Lic τ = 1.85 ±0.16 n=13). **(E)** Steady-state fast inactivation curves in G1475R variant Na^+^ channels in the presence of 300μM of S-Lic (blue) or vehicle (olive). Lines represent Boltzmann functions fit to the data points. G1475R variant induced a shift of the steady-state fast inactivation curve towards more depolarized potentials and exposure to 300μM of S-Lic did not alter those parameters. **(F)** Mean values of the recovery time constant at two different holding voltages (-80 and -100 mV, n=7 for G1475R, and n=8 for G1475R + 300 μM S-Lic). Inset of figure: Voltage-dependence of the major time constant of fast inactivation (n=10 for G1475R and for G1475R + 300 μM S-Lic). **(G)** Ramp current peaks (persistent current) normalized to transient peak currents I_peak_ upon ramp stimuli from -100 to +40 mV lasting 800 ms, n=11 for G1475R +/- S-Lic and n=8 for the others. **(H)** Peak transient Na^+^ currents normalized by cell capacitances (WT, n=14, WT + 300 μM S-Lic, n=9. M1760I, n=12, M1760I + 300 μM S-Lic, n=16, A1622D and A1622D + 300 μM of S-Lic, n=9, G1475R, n=10, G1475R + 300 μM of S-Lic, n=11. Shown are means ± SEM. WT (black), WT + 300 μM S-Lic (turquoise), M1760I (violet), M1760I + 300 μM S-Lic (pink), A1622D (red), and A1622D + 300 μM of S-Lic (orange), G1475R (olive green), G1475R + 300 μM of S-Lic (blue). Shown are means ±SEM for each data point. n, number of recorded cells; * p < 0.05; ** p < 0.01; *** p < 0.001; **** p < 0.0001 vs. vehicle (-S-Lic) respective group, the same goes for $ sign but indicates a difference vs. WT. Unpaired t-test or Mann-Whitney U-test were used for +/-S-Lic comparisons, for multiple group comparisons (WT+/-S-Lic and variant+/-S-Lic) one-way ANOVA with Dunnett's posthoc test or ANOVA on ranks with Dunn´s *posthoc* test were used.

**Supplementary Tables**

| **Transfected Na_V_1.6 channel** | **n** | **Current density at peak (pA/pF)** | **n** | **Steady-state activation** | | **n** | **τ_entry_ into slow inactivation (ms)** | **n** | **Steady-state slow inactivation** | | |
| --- | --- | --- | --- | --- | --- | --- | --- | --- | --- | --- | --- |
|  |  |  |  | **V_1/2_ (mV)** | **K** |  |  |  | **V_1/2_ (mV)** | **K** | **AUC** |
| **WT** | 14 | -343.1 ±49.78 | 10 | -20.14 ±1.54 | -6.06 ±0.48 | 10 | 3500 ±170.8 | 10 | -55.97 ±1.16 | 7.69 ±0.44 | 52.8 ±0.83 |
| **WT + 300uM S-Lic** | 9 | -232 ±35.63 | 9 | -20.11 ±1.08 | -5.59 ±0.38 | 11 | 2847 ±208.4 * | 7 | -72.06 ±0.71, **** | 6.05 ±0.54 * | 40.05 ±0.84 **** |
| **A1622D** | 9 | -349.4 ±65.49 | 9 | -10.07 ±1.11 $$$$ | -5.61 ±0.33 | 8 | 1423 ±93.3 $$$$ | 7 | -54.01 ±1.11 | 5.20 ±0.15 $$ | 52.8 ±1.27 |
| **A1622D +300uMS-Lic** | 9 | -216 ±29.03 | 9 | -6.60 ±0.95 $$$$, * | -6.18 ±0.34 | 10 | 871.1 ±53.58 $$$$, **** | 10 | -65.83 ±1.83 *** | 7.46 ±0.40 *** | 44.5 ±2.06 $$$, ** |
| **G1475R** | 10 | -268 ±44.87 | 10 | -16.83 ±1.76 | -5.50 ±0.49 | 13 | 3029 ±234.3 | 8 | -56.93 ±1.80 | 6.17 ±0.20 $ | 52.63 ±1.73 |
| **G1475R+300uMS-Lic** | 11 | -203.9 ±26.25 | 11 | -16.91 ±0.89 | -5.36 ±0.22 | 13 | 2182 ±202.4 $$$, * | 10 | -66.90 ±1.24 $$$$, *** | 5.11 ±0.21 $$$$, ** | 44.52 ±1.22 $$$, ** |
| **M1760I** | 12 | -478.6 ±71.8 | 12 | -26.10 ±1.58 $ | -4.79 ±0.37 | 12 | 4475 ±322.5 $ | 8 | -54.99 ±1.28 | 10.08 ±0.52 $$ | 55.01 ±1.15 |
| **M1760I+300uM S-Lic** | 16 | -464 ±52.82 | 16 | -27.95 ±1.42 $$ | -4.57 ±0.40 | 10 | 2984 ±355.4 ** | 10 | -70.66 ±1.35 $$$$, **** | 7.75 ±0.26 ** | 42.46 ±1.362 $$$$,**** |

**Table 1:** Data are presented as means ± SEM; n, number of recorded cells. AUC, area under the curve. * p < 0.05; ** p < 0.01; *** p < 0.001, **** p < 0.0001 vs. vehicle (-S-Lic) respective group, the same goes for $ sign but indicates a difference vs. WT. Unpaired *t*-test or Mann-Whitney U-test were used for +/-S-Lic comparisons, for multiple group comparisons (WT+/-S-Lic and variant+/-S-Lic) one-way ANOVA with Dunnett's *posthoc* test or ANOVA on ranks with Dunn´s *posthoc* test were used.

| **Transfected Na_V_1.6 channel** | **n** | **Steady-state fast inactivation** | | **n** | **Time course of fast inactivation at 0 mV (ms)** | **n** | **τ_rec_ (fast inactivation) at -100mV (ms)** | **n** | **Peak ramp current normalized to I_peak_** | **n** | **I_ss_/I_peak_**  **at 0 mV** |
| --- | --- | --- | --- | --- | --- | --- | --- | --- | --- | --- | --- |
|  |  | **V_1/2_ (mV)** | **K** |  |  |  |  |  |  |  |  |
| **WT** | 10 | -63.21 ±1.20 | 4.79 ±0.17 | 10 | 0.57 ±0.05 | 9 | 4.16 ±0.52 | 8 | -0.13 ±0.01 | 12 | 0.0129 ±0.0039 |
| **WT + 300uMS-Lic** | 7 | -66.98 ±1.08 * | 4.44 ±0.20 | 9 | 0.48 ±0.02 | 11 | 4.40 ±0.39 | 8 | -0.09 ±0.01 | 8 | 0.0111 ±0.0053 |
| **A1622D** | 7 | -56.31 ±1.41 $$$ | 7.76 ±0.18 $$$$ | 9 | 4.49 ±0.35 $$$ | 9 | 0.71 ±0.10 $$ | 8 | -0.25 ±0.03 $$ | 9 | 0.1135 ±0.0084 |
| **A1622D+300uMS-Lic** | 10 | -51.85 ±0.60 $$$$, ** | 8.40 ±0.37 $$$$ | 9 | 3.15 ±0.20 $, ** | 13 | 0.63 ±0.05 $$$ | 8 | -0.18 ±0.01 | 9 | 0.1093 ±0.0049 |
| **G1475R** | 8 | -58.04 ±0.87 $$ | 4.15 ±0.06 | 10 | 0.75 ±0.05 $ | 7 | 3.67 ±0.27 | 11 | -0.10 ±0.01 | 10 | 0.0120 ±0.0035 |
| **G1475R+300uMS-Lic** | 10 | -59.38 ±0.82 $ | 4.20 ±0.15 | 10 | 0.70 ±0.01 $ | 8 | 3.14 ±0.32 | 11 | -0.16 ±0.02 | 8 | 0.0158 ±0.0068 |
| **M1760I** | 8 | -65.04 ±0.90 | 4.63 ±0.24 | 10 | 0.67 ±0.03 | 7 | 3.95 ±0.35 | 8 | -0.12 ±0.01 | 11 | 0.0212 ±0.0015 |
| **M1760I+300uMS-Lic** | 10 | -65.77 ±1.35 | 4.80 ±0.18 | 10 | 0.52 ±0.02 ** | 7 | 2.79 ±0.14 $, * | 8 | -0.10 ±0.01 | 13 | 0.0196 ±0.0017 |

**Table 2:** Data are presented as means ± SEM; n, number of recorded cells. I_ss_, “steady-state‟ current. I_peak_, transient peak current. * p < 0.05; ** p < 0.01; *** p < 0.001, **** p < 0.0001 vs. vehicle (-S-Lic) respective group, the same goes for $ sign but indicates a difference vs. WT. Unpaired *t*-test or Mann-Whitney U-test were used for +/-S-Lic comparisons, for multiple comparisons (WT+/-S-Lic and variant+/-S-Lic) one-way ANOVA with Dunnett's *posthoc* test or ANOVA on ranks with Dunn´s *posthoc* test were used.
